# Short-Lived EEG Synchrony Patterns for Alzheimer’s Disease Diagnosis

**DOI:** 10.64898/2026.03.23.713571

**Authors:** Bilal Orkan Olcay

## Abstract

Developing a reliable detection of olfactory performance for early Alzheimer’s disease (AD) diagnosis remains challenging. Existing methods, such as psychophysical and event-related potential approaches, provide limited consistency in quantifying olfactory function. This study introduces a novel and objective framework that analyzes olfactory-stimulus-evoked EEG synchronizations of the subjects for AD diagnosis. We calculated the time-resolved wavelet coherence between EEG signals and then determined the timings (i.e., latency and duration) that describe when olfactory-stimulus-induced EEG channel interactions begin and end for each channel and frequency band. These timings, as well as the mean synchronization values in these segments, were used as features for diagnosis. Our framework, when cross-correntropy was used as a synchronization measure, exhibited a notable diagnostic accuracy in mild AD detection. The most discriminating feature between mild AD and healthy subjects was found to be the latency of synchronization between Fp1 and Fz in the low *θ* band, which showed significantly high correlation with clinical test scores. Furthermore, our framework achieved 100% diagnosis accuracy when EEG features and clinical test scores were used together. Our findings show that inter-channel short-lived synchronization timings serve as useful and complementary metrics about subjects’ olfactory performance and their neurological conditions.

## 1 Introduction

Alzheimer’s disease (AD) is the leading cause of dementia, marked by progressive cognitive and memory decline [1]. Early pathological changes arise in memory- and olfaction-related brain regions [2, 3]. However, symptom overlap with other dementias makes early diagnosis challenging [4]. Current clinical methods, including neuropsychological assessments and cerebrospinal fluid analysis, and neuroimaging techniques such as positron emission tomography (PET), single-photon emission computed tomography (SPECT), and magnetic resonance imaging (MRI), are largely effective only at advanced stages, limiting timely intervention [5]. Moreover, their applicability is limited due to their cost and implementation complexity. In that context, electroencephalography (EEG) is far more convenient for reaching more people for diagnosis. EEG signal analysis has been applied to various neurological disorders [6], with AD being one of the most prevalent [7, 8]. In the literature, extensive research has been conducted to evaluate EEG signals for diagnosing AD and their correlation with the outcomes of existing clinical approaches [9]. In these studies, entropy [10] and functional connectivity [11, 12, 13] appear as the most appealing information-theoretic methods for calculating informative features from a limited number of EEG sensors. Functional connectivity measures such as magnitude-squared and wavelet coherence, phase-locking index, and cross-correlation [14] have been used to unravel the alterations of brain interactions due to AD-related neurodegeneration [15]. The stationarity assumptions in the aforementioned functional connectivity-based studies limit the interpretability of the dynamically changing brain patterns. Upon this shortcoming, dynamic functional connectivity approaches have been put forward [13]. Findings from all these studies support the communication breakdown syndrome hypothesis, suggesting that AD pathology disrupts local neural interactions and progressively spreads to distant regions, and thus, impairs global brain synchrony and complexity patterns. Yet, the existing connectivity-based approaches may be affected by fluctuations in consciousness, wakefulness, and cognition during rest, reducing their accuracy and sensitivity to AD-related EEG patterns. To minimize such variations, memory and attention tasks are given to subjects to stabilize their physiological and psychological states [16].

Olfaction may be the earliest sensory domain to decline due to AD-related neurodegeneration, in which it starts many years before clinical symptoms appear [17]. In that context, the olfactory event-related potentials (OERP) and psychophysical olfactory tests have been used to diagnose AD subjects [17, 18]. The former approach measures the amplitudes and the latencies of averaged olfactory-evoked brain responses, while the latter measures odor identification, threshold, and discrimination capabilities of subjects. However, these approaches lack reliability since the OERP may not emerge for normosmic (healthy) participants or may even emerge for anosmic (people suffering from functional olfactory loss) participants [19]. Also, the latter approach (i.e., psychophysical olfactory performance test) is susceptible to the subjects’ response bias. Given these limitations of conventional methods, connectivity-based features from olfactory-evoked EEG offer a reliable and objective way to measure AD-related neural patterns. [12].

The olfactory system comprises a highly complex network in the human brain, extending from primary olfactory areas to broad cortical regions associated with emotional, semantic, learning, and memory processes without using thalamic relay [20, 21, 22]. Its anatomical projections extending from the olfactory to other structures explain why odor-based neural oscillations and their interactions are critical for monitoring neurodegenerative processes. Recent feature extraction methods in the literature, such as Wavelet-Spatial Domain Features, Sample Entropy, and Tunable Q-factor Wavelet Transform, have been used to analyze olfactory-stimulus-evoked EEG signals for diagnostic purposes [20, 23, 24]. However, many of the existing approaches rely on stationary assumptions or use predefined short-lasting time windows when calculating features from EEG channels, which overlook the brain activity’s dynamically changing nature. What is more, these studies exclude the fact that the brain is a complex network in which its regions exhibit reciprocal and intermittent interactions [25].

Despite their high accuracy, functional connectivity studies assume that the olfactory-evoked interactions among brain regions emerge and vanish simultaneously across all channel pairs and frequency bands. In addition, these studies do not conduct a detailed time-resolved brain connectivity/synchronization analysis to see when the olfactory-stimulus-evoked cortical interactions begin and how long they last. These two shortcomings, when considered together, means that existing approaches neglect the channel pair (*x, y*)- and frequency band (*f* )-specific latency 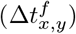 and duration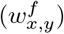timings of olfactory stimulus-evoked short-lived functional connectivity [12]. These timing parameters (i.e., synchrony latency and duration timings) are sensitive to the underlying neuronal activity [26], presumably altered due to AD neuropathology, making them substantial candidate features for detecting AD in mild stages. As evidence, olfactory stimulus-evoked short-lived brain connectivity patterns achieved remarkable Parkinson’s disease (PD) diagnosis accuracy [27]. Despite its novelty, [27] used a fixed-length time window and overlooked channel- and frequency-specific synchronization latency and duration in diagnosing PD.

In this study, we propose a novel and more comprehensive framework for AD diagnosis purposes. Our framework calculates time-resolved brain connectivity patterns and obtains the latency and duration timings of olfactory-stimulus evoked short-lived brain synchronization as features. Thus, our framework considers the fact that stimulus-evoked brain interactions for each channel pair and frequency band emerge for a certain period of time after olfactory stimulus onset, which is neglected by studies adopting stationarity of the functional connectivity. We investigate (1) whether these olfactory-stimulus-evoked brain synchronization timings are systematically altered in individuals with mild AD compared to healthy control (HC) subjects, and (2) whether these timings, together with statistical features derived from them, can provide notable AD diagnosis performance. For evaluation, we used a publicly available olfactory EEG dataset [28].

## 2 Materials and Methods

### 2.1 Dataset and Synchronization Measures

We used an online-available Olfactory EEG dataset in [28] to assess the diagnosis accuracy of our framework. This dataset comprises the olfactory stimulus-induced EEG signals collected from 13 HC and 11 mild AD patients while they were given either lemon or rose odorant stimuli in a random fashion. The lemon odor was the frequent odorant (75%), and the rose was the rare (25%). Approximately 120 stimuli were presented to each subject using the olfactometer setup proposed in [29]. Throughout the study, we used the EEG periods in which the lemon odorant was presented to subjects [30]. The mean ages of mild AD subjects and HC are 76.63 *±* 9.22 and 68.23 *±* 6.18, respectively.

The EEG signals were collected via a 32-channel EEG recording system with a sampling frequency of 2 kHz. The EEGLAB toolbox was used for preprocessing the EEG signals before public distribution [31]. The preprocessing steps include 0.5–40.5 Hz band-pass filtering and ICA-based artifact removal. Additionally, a procedure was used to remove artifact-contaminated stimulus periods from further analyses. The publicly shared version of this dataset includes four-channel (Fp1, Fz, Cz, Pz) preprocessed EEG signals of olfactory stimulus periods. The online-accessible preprocessed EEG signals, which we downloaded and used, were down-sampled to a sampling frequency of 200 Hz before being shared online. Each stimulus period begins 1 second before the stimulus onset and continues until 2 seconds after.

In this study, we calculated coherences between EEG channels for each time instant and frequency band using wavelet scalograms using all 3-second-long stimulus periods, similar to the wavelet coherence calculation approach in [32]. The six different synchronization measures we used to calculate the synchronization of scalograms are cross-covariance, Kraskov’s mutual information estimation, Kendall’s tau correlation, cross-correntropy, cosine similarity, and nonlinear interdependency measure [33]. The mathematical expression of each synchronization measure is provided as Supplementary Materials using the scalogram variables 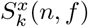 and 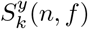obtained from channels *x* and *y* for a particular time instant *n* and frequency band *f* .

### 2.2 Proposed Method

The operational flow of the proposed measurement framework is depicted in Fig. 1, while Fig. 2 presents wavelet scalograms of four EEG electrodes for an HC subject and an AD subject. In the *Wavelet Transform* step, wavelet coefficients of all EEG channels of each stimulus period were calculated using a complex Morlet wavelet function from 1 Hz to 40.5 Hz with 0.5 Hz steps. During calculation, we adopted the procedure in [34]. The mean wavelet scalograms at each time instant *n* for a frequency band *f* could be calculated as:

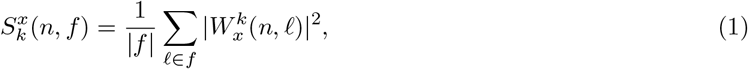

where 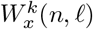denotes the wavelet coefficient calculated at time instant *n* and frequency point *𝓁* for channel *x* and stimulus period *k*. |*f*| gives the total number of frequency points within the frequency band *f* . Here, we divided each conventional frequency band (i.e., *δ, θ, α, β*, and *γ*) into two equal pieces and obtained ten frequency sub-bands as given in Table 1.

**Figure 1:**
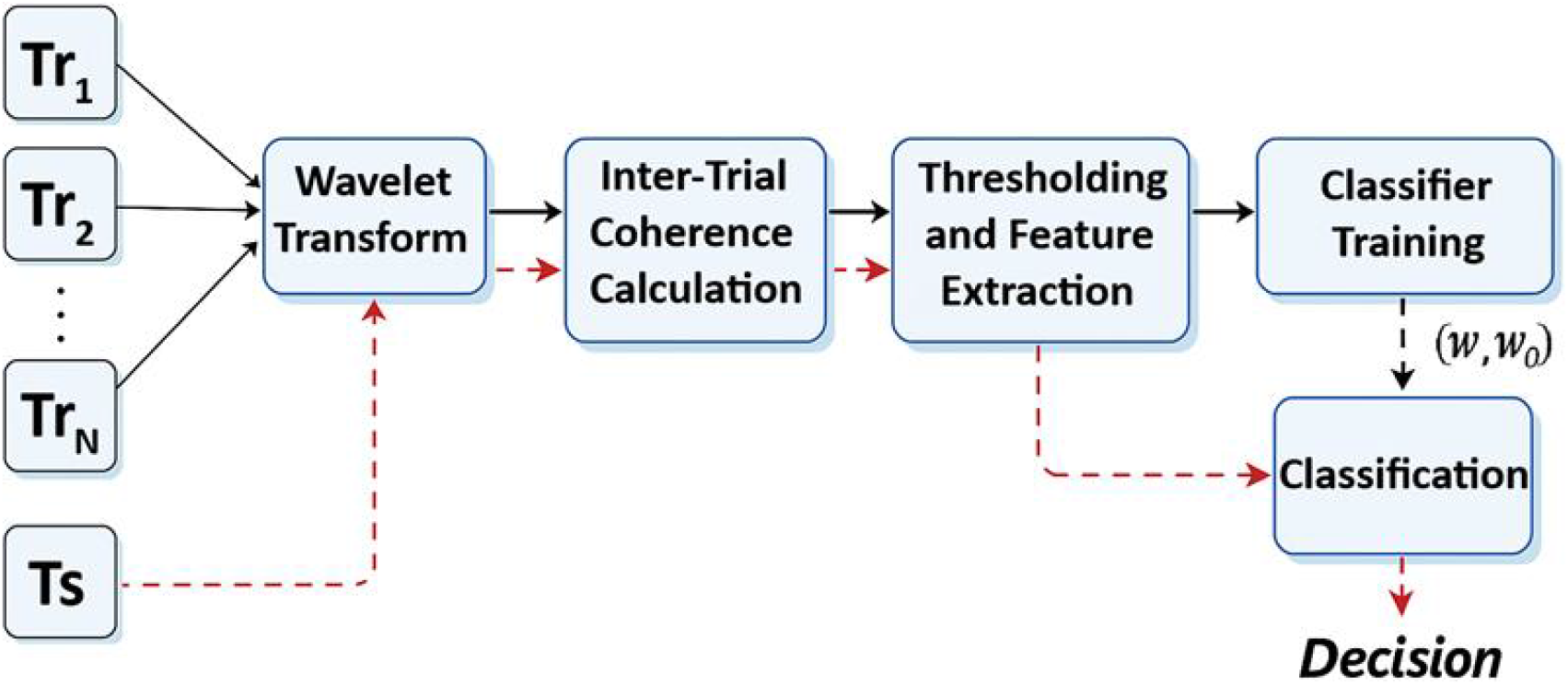
The operational flow diagram of the proposed measurement framework. Tr_1_, Tr_2_, …, Tr_*N*_ denote training subjects, and Ts denotes the unseen test subject.

**Figure 2:**
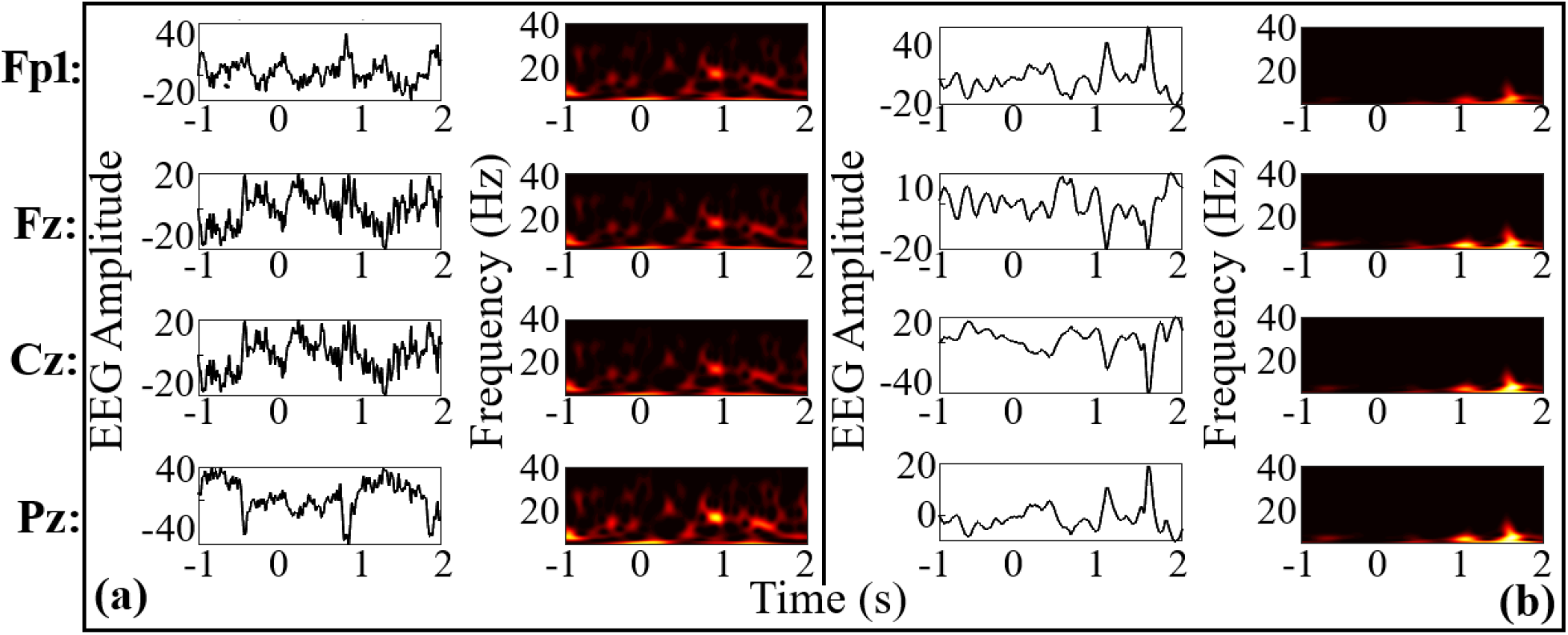
An illustration of EEG activities of four electrodes and their complex Morlet wavelet scalograms of (a) a healthy control subject, (b) a subject with mild Alzheimer’s disease.

**Table 1:** Low and High Sub-band Frequency Ranges for EEG Signals.

| Sub-band | $\delta$ | $\theta$ | $\alpha$ | $\beta$ | $\gamma$ |
| --- | --- | --- | --- | --- | --- |
| Low [Hz] | [1, 2] | [4, 5.5] | [8, 10] | [13, 21.5] | [31, 35.5] |
| High [Hz] | [2.5, 3.5] | [6, 7.5] | [10.5, 12.5] | [22, 30.5] | [36, 40.5] |

In the *Inter-Trial Coherence Calculation* step, we calculated the coherence 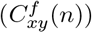between scalogram coefficients of channels *x* and *y* using all the stimulus periods indexed by *k* for each frequency band *f* and time instant *n* as [32]:

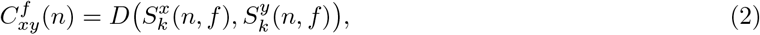

where *D*(·, ·) denotes the synchronization measure of choice.

In the *Thresholding and Feature Extraction* step, we first determined a threshold for each frequency band and channel pair through the Kolmogorov-Smirnov (KS) test. Briefly, the threshold was defined as the coherence value yielding the maximum distance between the prestimulus ([*−*400ms, 0ms]) and poststimulus ([0s, 2s]) CDFs [35]:

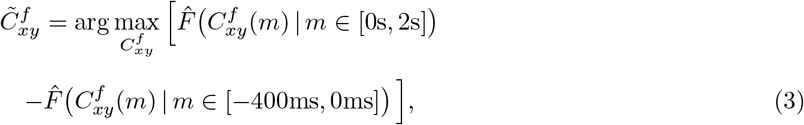

where 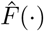provides the empirical cumulative distribution of coherence values.

We utilize the [*−*400ms, 0ms] time interval to represent the electrophysiological behavior of the brain before stimulus onset [36]. We used the KS test rather than a conventional hypothesis test to determine the coherence threshold. For each subject, channel pair, and frequency band, the threshold was set to the coherence value yielding the maximum divergence between prestimulus and poststimulus cumulative distributions of instantaneous coherences, providing an objective and non-parametric separation of baseline and stimulus-evoked interactions. Olfactory-related short-lived synchronizations were then determined by comparing stimulus-locked coherence values at each time instant exceeding this threshold. Next, the longest coherence segment surviving after thresholding was selected from 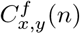as the dominant olfactory-related one, and the remaining shorter segments were discarded. We then used the latency and duration of this longest segment, and also the mean of coherence within this segment, as features. These three features were extracted from each channel pair and frequency band, yielding a total of 180 features per subject (6 channel pairs *×* 10 bands *×* 3 features).

In the *Classifier Training* step, we used the training subjects’ feature vectors to calculate the Fisher score of features to obtain a discriminative feature subset of far fewer than 180 features. To that end, we selected the features having a score greater than the mean plus two times the standard deviation of all Fisher scores. These selected features were then used for classifier training. Please note that the Fisher-score–based feature selection was performed only on the training subjects within each cross-validation cycle. At each cross-validation cycle, we selected a useful feature subset as described above and used it for training the classifier for diagnosis purposes. Four classifiers were evaluated: Fisher’s linear discriminant (FLD), linear and nonlinear support vector machines (SVM), and *k*-nearest neighbor (kNN), all well established in machine learning. As regards the nonlinear SVM, the feature vectors are transformed to another space via a kernel function as in [37]. The kernel function of nonlinear SVM was selected as radial basis function whose kernel width parameter *ρ* was calculated as [33]:

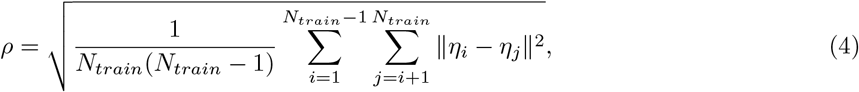

where *η*_*i*_ is the feature vector obtained from training subject *i*, and *N*_*train*_ is the number of subjects in the training set.

Finally, for the kNN classification algorithm, the *k* parameter was set to 5 and the Manhattan distance was used [38], resulting in optimal classification results.

In the test phase (red dashed lines in Fig. 1), all steps are identical to those in the training phase except classifier training. During the *Classification* step, the trained classifier with reduced feature set in the training set was used to decide the category of the unseen test subject’s reduced feature vector. To evaluate the recognition performance of the proposed measurement framework, a leave-one-subject-out cross-validation scheme was adopted, where one subject was reserved for testing and the remaining ones for training. The cross-validation cycles were repeated until each subject was excluded once for testing purposes. The average performance is calculated as the mean of the performance attained at every cross-validation cycle. The steps are outlined in Algorithm **??**.

The AD subjects were statistically significantly older than the HC, which could introduce age-related bias in the EEG features [39]. To mitigate this potential confounding effect, an age-detrending procedure was incorporated into the analysis pipeline. At each cross-validation cycle, the Spearman rank correlation between each EEG feature and age was evaluated using the healthy training subjects only. Features exhibiting a near-significant association with age (p ≤ 0.1) were detrended using a generalized linear model (GLM) [30]. Specifically, for each corresponding cross-validation fold, a GLM with a Gaussian family and identity link was fitted using training subjects only, modeling the selected EEG feature as a linear function of age. The simple equation for the GLM model we adopted here is *Y* = *β*_0_ + (*β*_1_ *× X*_*age*_) + *ϵ*. The residuals (i.e., *Y − Y*_*estimated*_) of this model were then used as age-detrended features for both training and test subjects within the same fold. This procedure was applied solely as a preprocessing step to remove linear age-related trends.

## 3 Results

### 3.1 Classification Performances and Biophysical Results

The overall performances of our framework in classifying AD versus HC subjects for different synchronization measures are given in Fig. 3. Our framework achieved 87.50% accuracy when cross-correntropy measure was used. Besides, Kraskov’s mutual information and Kendall’s tau correlation methods achieved 83.33% and 75% accuracies, respectively. However, cross-covariance, cosine similarity, and nonlinear interdependency measures did not elicit any marked performances. Synchronization measures achieving over 70% accuracy were mostly nonlinear. While cross-covariance and cosine similarity capture only second-order statistical interactions, nonlinear interdependency exploits Euclidean distances between phase-space vectors in scalogram synchronization. These results indicate that nonlinear measures are more effective in revealing olfactory-evoked brain synchronizations than those limited to second-order interactions.

**Figure 3:**
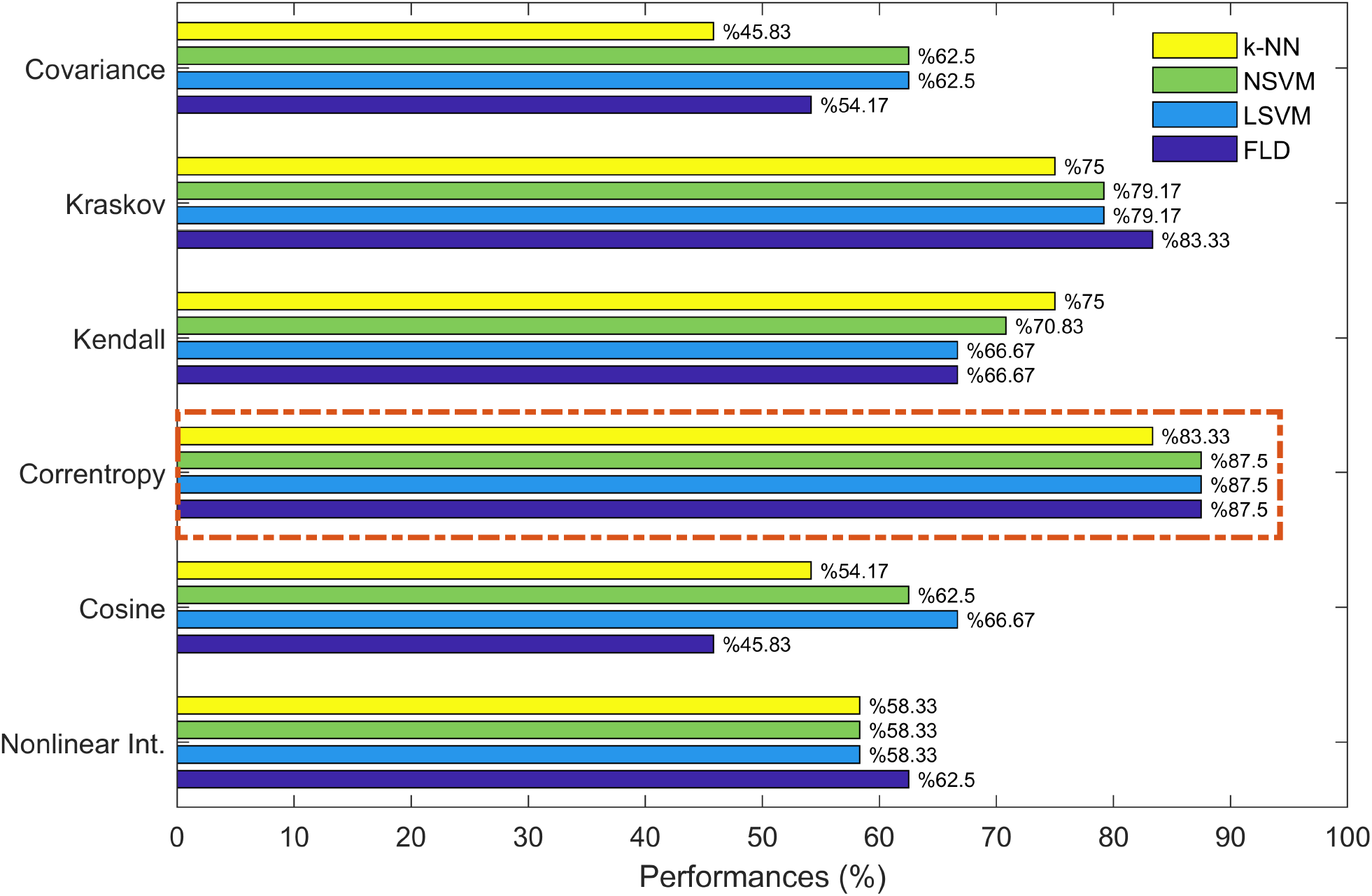
Average diagnosis accuracy performance for different synch. measures used in the proposed framework.

We investigated the frequently selected features across cross-validation cycles when the correntropy method was used in our framework. Only the latency of Fp1–Fz synchronization at low *θ* band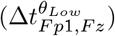was selected and used at all cross-validation cycles. On the contrary, none of the other features were selected within cross-validation cycles, meaning that the features except 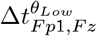did not stably reflect discriminative information related to AD neurodegeneration.

We demonstrated the distributions of 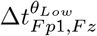 feature for AD and HC groups in Fig. 4. According to distributions, stimulus-evoked synchronization latency of channels Fp1–Fz at low theta band are significantly different for subjects with mild AD and HC. In addition, we quantified the distinctiveness of all features by *t*-test analysis. We corrected the resulting *p*-values against multiple comparisons via the Benjamini–Hochberg procedure. The corrected *p*-value results revealed that only the Fp1–Fz synchronization latency at low theta was statistically significantly different for these two groups (*p*_*corrected*_ = 0.014). In addition to the *p*-values, we also calculated the statistical power of the latency feature 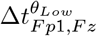using Hedges’ *g*-test. The effect size was calculated as follows: Cohen’s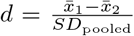, and the correction factor was applied to derive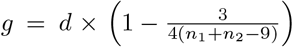. Our analysis showed an extremely large effect size of *g* = 1.87, confirming the robustness of the observed effects despite the small sample size. Please note that the *p*-values of all age-detrended latency features are provided in Supplementary Materials.

**Figure 4:**
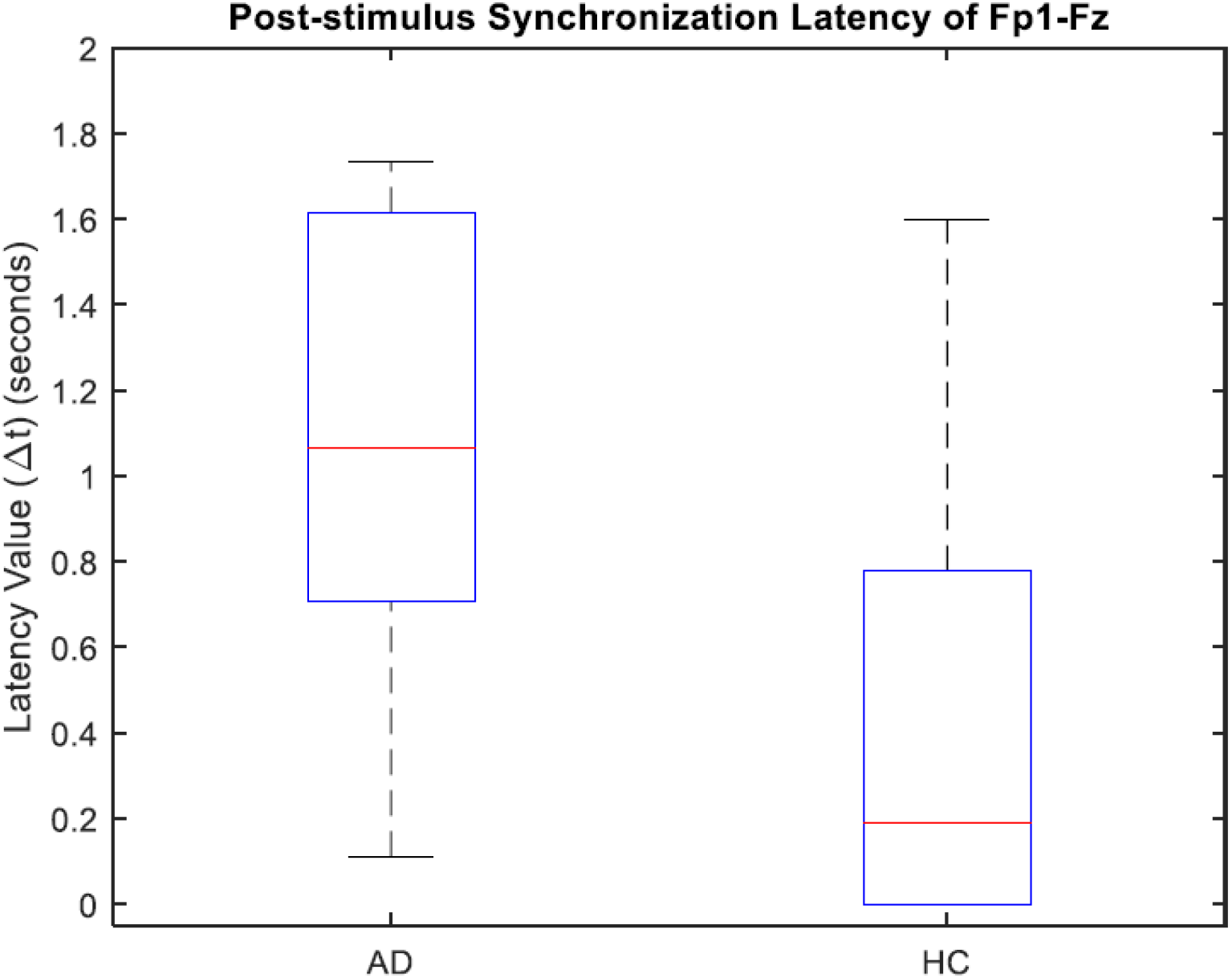
The distribution of synchronization latency of Fp1–Fz channel pair within low theta(*θ*_*Low*_) frequency band 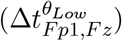for both AD and HC groups.

We also calculated the Spearman rank correlation of 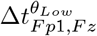feature with clinical University of Pennsylvania Smell Identification Test (UPSIT) and Mini-Mental State Examination (MMSE) test scores. We observed that 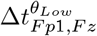exhibited strong and statistically significant correlations with UPSIT (*ρ*_*UP SIT*_ = *−*0.552, *p* = 0.0052) and MMSE (*ρ*_*MMSE*_ = *−*0.531, *p* = 0.007) scores. These results indicate a negative correlation between *θ*^*Low*^-band frontal synchronization latency and clinical test scores: as AD progresses, Fp1–Fz latency increases, reflecting a decline in clinical performance. This proves that our EEG-based olfactory measure aligns well with established clinical scores.

Age-detrended feature variations are illustrated in Fig. 5. As shown in Fig. 5, only one AD subject exhibited a negative feature value, whereas the remaining AD subjects showed positive values. Similarly, for the HC subjects, while only two of them elicited positive-valued features, the remaining HC subjects had negative-valued ones. These findings show that 21 out of 24 subjects exhibited latency features aligned with their respective groups, meaning that the latency of synchronization between Fp1 and Fz channels in the *θ*^*Low*^ band could classify the subjects with an 87.50% success rate.

**Figure 5:**
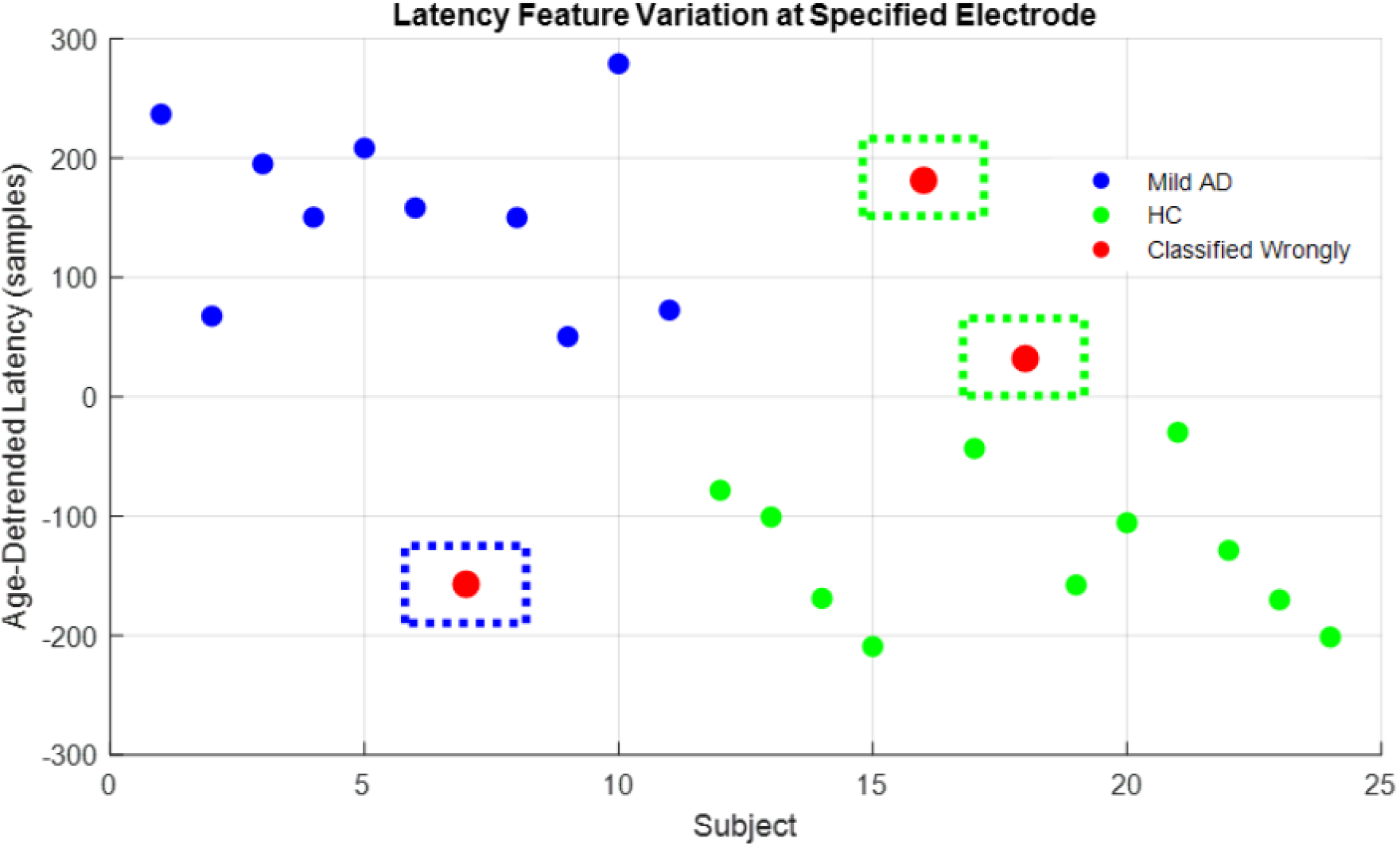
The latency feature 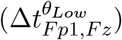variation. The latency feature values shown are age-detrended. Red samples enclosed by blue and green rectangles represent subjects whose latency values deviate from their respective groups.

To further assess the contribution of synchronization timing parameters, an ablation analysis was performed by excluding the synchronization latency and considering only the synchronization duration as features. We determined and used only the duration parameter that provides maximal deviation between pre- and post-stimulus cumulative instantaneous distributions. Under this setting, the proposed framework achieved a classification accuracy of 62.50%, which is substantially lower than the performance obtained when latency information was incorporated. This marked reduction in accuracy demonstrates that inter-regional synchronization latency constitutes a critical characteristic that changes upon neurodegeneration.

Table 2 also presents the features identified by other synchronization measures achieving over 70% accuracy. Each different synchronization measure put forward different features from distinct frequency bands and channel pairs. The observed variability in synchronization features across synchronization measures presumably stems from their differing sensitivities to neural dynamics [40, 41]. Specifically, proposed method frequently selected and used duration and mean of coherence features generally from the Fp1–Pz channel pairs when using Kraskov’s method. Besides, the latency and duration parameters were calculated from the channel pairs constructed with Fp1 when the Kendall tau correlation method was used. Although the selected and used features across cross-validation cycles are specific to the synchronization measure used throughout the analysis, most features providing more than 70% diagnosis accuracy were derived from the channel pairs constructed with the frontal Fp1 channel.

**Table 2:**
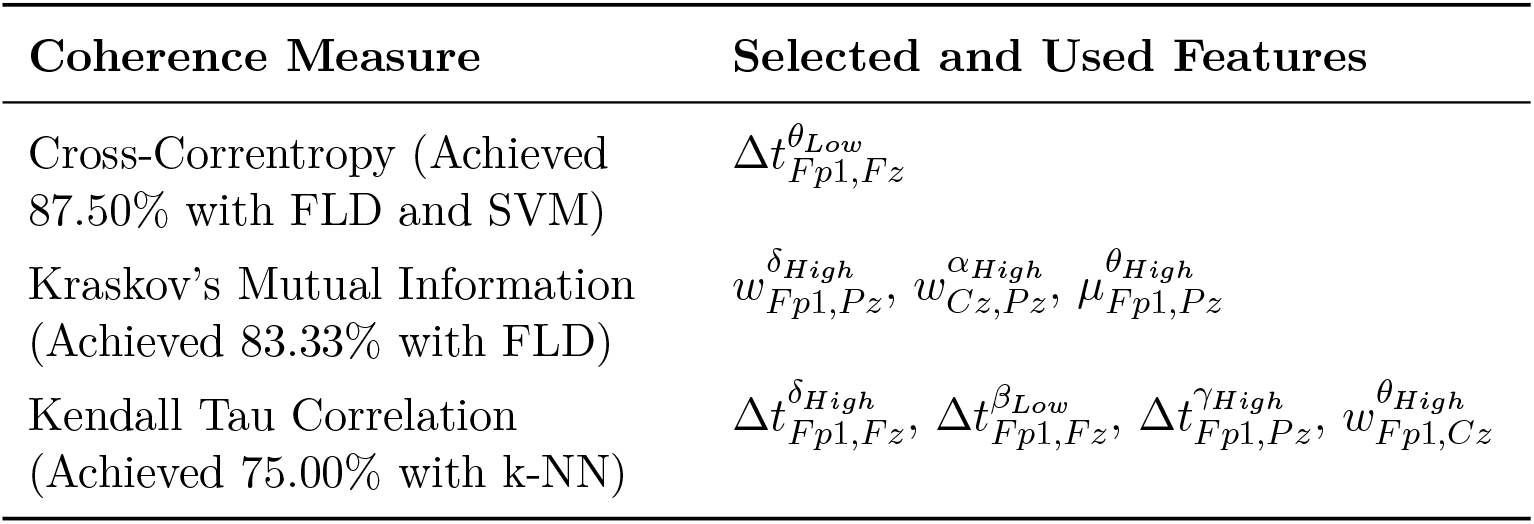
Frequently selected features by synchronization methods whose performance exceeded 70%.

### 3.2 Performance Comparisons with Existing Methods

Table **??** summarizes the key features of methods reported in the literature. While psychophysical test results, such as the UPSIT and threshold-discrimination-identification (TDI) scores, rely heavily on subjective responses and therefore lack objectivity, EEG-based electrophysiological approaches provide quantitative and expert-free measurements. However, previous EEG studies have generally focused on amplitude or latency properties (e.g., OERP) or static connectivity indices (e.g., imaginary coherence), which do not tackle the timings of short-lived neural synchronizations [27, 30, 36]. In contrast, the proposed framework provides a fully objective and temporally-detailed EEG characteristics using synchronization timings without requiring any behavioral input from the participant.

We compared our proposed measurement framework with five existing EEG-based methods. The first uses imaginary coherences of Fz–Cz channel pairs in the *β* and *γ* bands [30]. The second calculates EEG entropies in Cz and Pz channels over three short windows—pre-[-400 ms, 0 ms], during-[400 ms, 800 ms], and post-stimulus [1100 ms, 1500 ms]—after filtering (0.5–30 Hz), averaging first across channels and then across stimulus periods to obtain subject-specific three-dimensional feature vectors [36]. The third benchmark method calculates and uses the mean olfactory-stimulus-evoked event-related spectral perturbation (ERSP) using 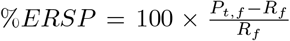 formula within the frequency range of [2, 6]*Hz* and the time window of [200, 2000]*ms* following stimulus onset [42]. Here, *P*_*t,f*_ represents the mean wavelet scalogram value at time *t* and frequency *f*, and *R*_*f*_ denotes the mean scalogram across the prestimulus interval at frequency *f* . The results in Table 4 indicate that the proposed method outperforms the entropy-based, coherence-based, and *t*–*f* ERSP benchmarks. Furthermore, recent advancements utilizing sophisticated signal decomposition and entropy-based deep learning models have reported high classification accuracies, yet these approaches often prioritize predictive performance over the direct physiological interpretability of the underlying neural mechanisms [23, 24]. In contrast, our proposed measurement framework focuses on explicit, data-driven synchronization timings that remain highly discriminative even under simpler classification schemes and rigorous age-detrending procedures. The attainment of 100.00% diagnostic accuracy when these features are integrated with clinical scores further suggests that time-resolved synchronization parameters provide a fundamental and objective basis for assessment that effectively complements traditional neuropsychological evaluations.

**Table 3:** Subject diagnosis performance comparison of benchmark methods.

| Method | EEG Features (%) | EEG + UPSIT (%) |
| --- | --- | --- |
| Proposed Method (Correntropy-based) | **87.50** | **100.00** |
| Ref. [30] (ImCoh of Fz–Cz) | 83.30 | 91.70 |
| Ref. [36] (K–L Entropy, Three Windows) | 58.33 | 70.83 |
| Ref. [42] (EEG Time–Frequency Power Change) | 62.50 | 87.50 |
| Ref. [23] (CSP, CMTS, TQWCM) | 93.14 | N.A. |
| Ref. [24] (Sample Entropy) | 92.14 | N.A. |

**Table 4:**
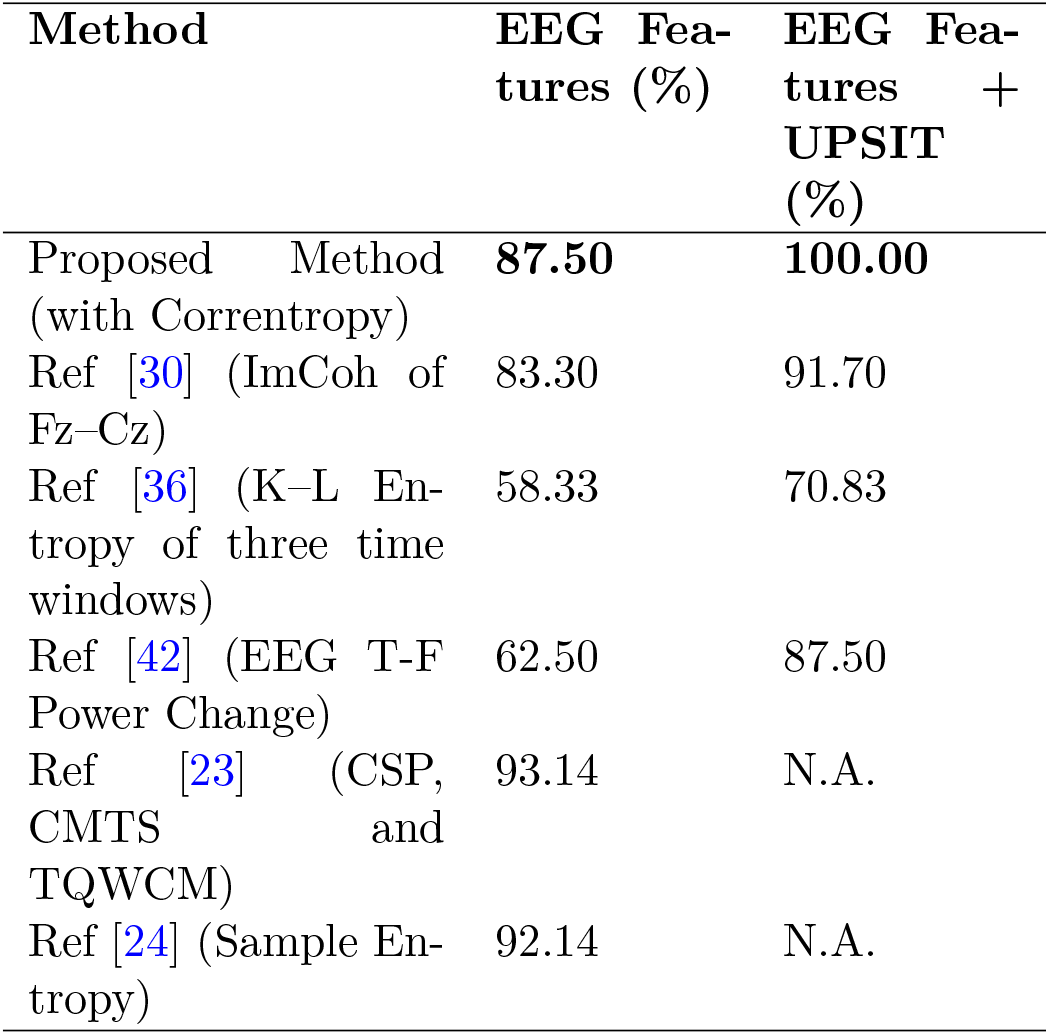
The subject diagnosis performances of the methods used for benchmark comparisons.

## 4 Discussion

The motivation for proposing this method is as follows: AD-related disturbances alter neural activity characteristics across various states, such as resting, sensory, motor, and perceptual processes [5]. Among these, olfactory deficits emerge as one of the earliest to be degenerated in AD. Deficits in this system are a result of several neurodegenerations, like the deposition of amyloid plaques and the loss of synapses within olfactory-related subcortical (such as hippocampus, entorhinal, and piriform cortex) and cortical structures [17, 43], which inhibits, delays, and deteriorates their sensory information processing. This study aims to characterize these neural alterations by investigating 1) the latency, 2) the duration, and 3) the strength (mean) synchronization of this short-lived synchronous activity for AD diagnosis purposes. Unlike previous olfactory-stimulus-induced EEG studies that primarily focused on event-related potential (ERP) amplitudes or on the statistical interdependence of fixed-length EEG segments, our approach offers a new methodology for examining the temporal organization of short-lasting inter-regional synchronization of cortical activity as a biomarker of AD-related neurodegeneration.

In our framework, the latency feature 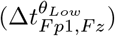ranged from 0–780 ms for HC and 750–1600 ms for Alzheimer’s patients (see Fig. 4). An increased synchronization latency was observed between the Fp1–Fz channel pair in the low-*θ* band 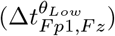in AD subjects in which this feature was consistently selected across all leave-one-out cross-validation cycles. Furthermore, this latency parameter elicited significant negative correlations with UPSIT and MMSE scores. This clearly indicates that this timing parameter is a stable AD biomarker and reflects AD-related cognitive and olfactory decline robustly.

The latency results obtained here for HC subjects are in line with existing literature reporting that initial cortical decoding and perceptual processing of odors typically lasts approximately 600 ms post-stimulus [20]. In contrast, the AD latency results show that frontal synchronization in the *θ* frequency band is markedly delayed. Given that *θ* activity reflects odor content processing [44], decision-making, and memory retrieval through hippocampal interactions [45, 46], and that the orbitofrontal cortex serves as a convergent area for olfactory information [47], these delays are likely due to impaired temporal coherence across these regions. Ultimately, such delays in frontal *θ* synchronization timing manifest as the characteristic olfactory and cognitive deficits observed in Alzheimer’s disease.

### 4.1 Limitations of the Study

In this study, we used an online available dataset with a cohort size of 24, which may raise several generalizability and overfitting questions. To mitigate these confusions, we first adopted a leave-one-out cross-validation procedure to validate the effectiveness of our approach. Second, the Hedges’ *g* effect size calculated in the study is an extremely high value of 1.87, which proves that the observed performance and biophysical are not coincidental and have a biological basis, even though the sample size is small. The scarcity of open-access olfactory EEG datasets involving clinical Alzheimer’s populations underscores the necessity of utilizing standardized benchmark data for the initial validation of novel methodologies.

Another limitation is that we used only 4 EEG sensors (Fp1, Fz, Cz, Pz) data due to limitations in this online Olfactory EEG dataset. Although this seems as a limitation, in several clinics, portable EEG devices with a low number of channels or wireless sensors are preferred to overcome size and battery lifetime issues. Existing studies aimed to achieve high performance with fewer channels [48], which is also essential for making our approach practical for clinical use. The fact that only the same feature is obtained repeatedly due to the limited number of EEG channels may be seen as a drawback. Yet, the consistent selection of only Fp1-Fz *θ* latency feature is not a weakness, but rather is a proof of its strength and parsimoniousness.

The fact that a single feature among 180 features remains stable across all cycles indicates that this feature is disease-specific and unaffected by noise or subject differences. The statistically significant (*p ≤* 0.01) and strong correlations of this single feature with clinical UPSIT and MMSE test scores exhibit its biophysical validity.

In this dataset, the AD and HC groups show a statistically significant age difference that many of the existing studies neglect. This age difference may appear in EEG features and may cause classification performance higher than it should be. To mitigate this confounding effect, in each cross-validation cycle, the relationship between age and EEG features was calculated with Spearman correlation, and those found to be significantly correlated were age-detrended by using a linear model. All the classification and analysis conducted here used age-detrended EEG features rather than the originally extracted ones. This means that the performances obtained here are due to disease, not age.

We obtained the timing parameter features with a 5-millisecond resolution (*f*_*s*_ = 200 Hz). Using an EEG dataset that is collected with a higher sampling frequency may provide a precise synchronization timing parameter determination and thus accurate brain activity characterization.

## 5 Conclusion

We proposed a novel framework that captures and uses as features of the short-lived EEG channel synchronization initiation and duration timings upon olfactory stimuli. Our results exhibit that the latency of frontal low-*θ* synchronization (i.e., 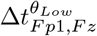) emerged as the most discriminative feature, showing strong correlations with UPSIT and MMSE scores. These findings indicate that delayed synchronization reflects Alzheimer ‘s-related neurodegeneration and highlight the potential of time-resolved EEG synchronization as an early and objective biomarker.

## Acknowledgments

We would like to thank Anil Karatay and Enes Ataç for their valuable contributions to the preparation of this paper.

