## Supplementary File-1 for "Short-Lived EEG Synchrony Patterns for Alzheimer’s Disease Diagnosis"

### Supplementary Materials

#### 1 Synchronization Measures

##### 1.1 Cross-Covariance

The cross-covariance between variables $S^x(n, f)$ and $S^y(n, f)$ is defined as [1]:

$$D_{\text{cov}}(S^x, S^y) = \frac{1}{N_k - 1} \sum_{k=1}^{N_k} \left(S_k^x - \mu_{S^x}\right)\left(S_k^y - \mu_{S^y}\right), \tag{1}$$

where $S_k^x(n, f)$ is the scalogram of the $k$-th trial and $\mu_{S^x}$ is the mean across all trials.

##### 1.2 Kraskov's Mutual Information

Kraskov's estimator calculates mutual information when joint PDFs cannot be evaluated [2]:

$$D_{\text{MI}}(S^x, S^y) = \psi(k) - \langle \psi(n_{S^x} + 1) + \psi(n_{S^y} + 1) \rangle + \psi(N_k), \tag{2}$$

where $\psi(\cdot)$ is the digamma function and $n_{S^x}$, $n_{S^y}$ denote neighbor counts within distance $\frac{\epsilon(i)}{2}$ [3].

##### 1.3 Kendall's Tau Correlation

Kendall's rank correlation is defined as [1, 4]:

$$D_{\text{Kendall}}(S^x, S^y) = \frac{2\xi}{N_k(N_k - 1)}, \tag{3}$$

with

$$\xi = \sum_{i=1}^{N_k - 1} \sum_{j=i+1}^{N_k} \text{sign}\left[\left(S_i^x - S_j^x\right)\left(S_i^y - S_j^y\right)\right]. \tag{4}$$

#### 1.4 Cross-Correntropy

Correntropy measures nonlinear statistical dependence by considering time and probability structure. The mathematical formulation is given as [5,6]

$$D_{\text{correntropy}}(S^x, S^y) = \frac{1}{N_k} \sum_{k=1}^{N_k} \kappa\big(S_k^x,\, S_k^y\big), \tag{5}$$

where the Laplacian kernel is defined as $\kappa(z_1, z_2) = \exp(-|z_1 - z_2|)$ [7].

#### 1.5 Cosine Similarity

Cosine similarity between $S^x(n, f)$ and $S^y(n, f)$ is [8]:

$$D_{\text{cosine}}(S^x, S^y) = \pi - \arccos\left( \frac{\sum_{k=1}^{N_k} S_k^x S_k^y}{\sqrt{\sum_{k=1}^{N_k} (S_k^x)^2}\,\sqrt{\sum_{k=1}^{N_k} (S_k^y)^2}} \right). \tag{6}$$

#### 1.6 Nonlinear Interdependency

Phase-space vectors are constructed as [9]:

$$\mathbf{S}_i^x = (S_i^x, S_{i-d}^x, \ldots, S_{i-(m-1)d}^x)^T, \tag{7}$$

$$\mathbf{S}_i^y = (S_i^y, S_{i-d}^y, \ldots, S_{i-(m-1)d}^y)^T. \tag{8}$$

The average distance to the first $k$ neighbors is:

$$R_n^k(S^x) = \frac{1}{k} \sum_{\ell=1}^{k} \|\mathbf{S}_n^x - \mathbf{S}_{\Gamma_n(\ell)}^x\|^2, \tag{9}$$

and the conditional distance:

$$R_n^k(S^x \mid S^y) = \frac{1}{k} \sum_{\ell=1}^{k} \|\mathbf{S}_n^x - \mathbf{S}_{\zeta_n(\ell)}^x\|^2. \tag{10}$$

Finally, the nonlinear interdependency is:

$$D_{\text{nonlinearInt}}(S^x, S^y) = \frac{1}{N} \sum_{n=1}^{N} \frac{R_n^k(S^x)}{R_n^k(S^x \mid S^y)}, \tag{11}$$

where $k = 6$ and $d = 10$ follow [10,11].
