## Supplementary File-2 for "Short-Lived EEG Synchrony Patterns for Alzheimer’s Disease Diagnosis"

| No. | Frequency Band | Channel Pair | Feature Type | P-Value (B-H Corrected) | Significance |
|---|---|---|---|---|---|
| 1. | low delta | Fp1-Fz | Latency | 0,999432384 | Not Significant |
| 2. | high delta | Fp1-Fz | Latency | 0,462612914 | Not Significant |
| 3. | low theta | Fp1-Fz | Latency | 0,014391469 | Significant |
| 4. | high theta | Fp1-Fz | Latency | 0,462612914 | Not Significant |
| 5. | low alpha | Fp1-Fz | Latency | 0,975742021 | Not Significant |
| 6. | high alpha | Fp1-Fz | Latency | 0,975742021 | Not Significant |
| 7. | low beta | Fp1-Fz | Latency | 0,975742021 | Not Significant |
| 8. | high beta | Fp1-Fz | Latency | 0,975742021 | Not Significant |
| 9. | low gamma | Fp1-Fz | Latency | 0,998362565 | Not Significant |
| 10. | high gamma | Fp1-Fz | Latency | 0,975742021 | Not Significant |
| 11. | low delta | Fp1-Cz | Latency | 0,975742021 | Not Significant |
| 12. | high delta | Fp1-Cz | Latency | 0,975742021 | Not Significant |
| 13. | low theta | Fp1-Cz | Latency | 0,975742021 | Not Significant |
| 14. | high theta | Fp1-Cz | Latency | 0,975742021 | Not Significant |
| 15. | low alpha | Fp1-Cz | Latency | 0,998362565 | Not Significant |
| 16. | high alpha | Fp1-Cz | Latency | 0,975742021 | Not Significant |
| 17. | low beta | Fp1-Cz | Latency | 0,999432384 | Not Significant |
| 18. | high beta | Fp1-Cz | Latency | 0,975742021 | Not Significant |
| 19. | low gamma | Fp1-Cz | Latency | 0,975742021 | Not Significant |
| 20. | high gamma | Fp1-Cz | Latency | 0,987853791 | Not Significant |
| 21. | low delta | Fp1-Pz | Latency | 1 | Not Significant |
| 22. | high delta | Fp1-Pz | Latency | 0,975742021 | Not Significant |
| 23. | low theta | Fp1-Pz | Latency | 0,975742021 | Not Significant |
| 24. | high theta | Fp1-Pz | Latency | 0,975742021 | Not Significant |
| 25. | low alpha | Fp1-Pz | Latency | 0,975742021 | Not Significant |
| 26. | high alpha | Fp1-Pz | Latency | 0,975742021 | Not Significant |
| 27. | low beta | Fp1-Pz | Latency | 0,462612914 | Not Significant |
| 28. | high beta | Fp1-Pz | Latency | 0,462612914 | Not Significant |
| 29. | low gamma | Fp1-Pz | Latency | 0,975742021 | Not Significant |
| 30. | high gamma | Fp1-Pz | Latency | 0,975742021 | Not Significant |
| 31. | low delta | Fz-Cz | Latency | 0,999432384 | Not Significant |
| 32. | high delta | Fz-Cz | Latency | 0,987853791 | Not Significant |
| 33. | low theta | Fz-Cz | Latency | 0,975742021 | Not Significant |
| 34. | high theta | Fz-Cz | Latency | 0,975742021 | Not Significant |
| 35. | low alpha | Fz-Cz | Latency | 0,998362565 | Not Significant |
| 36. | high alpha | Fz-Cz | Latency | 0,975742021 | Not Significant |
| 37. | low beta | Fz-Cz | Latency | 0,975742021 | Not Significant |
| 38. | high beta | Fz-Cz | Latency | 0,998362565 | Not Significant |
| 39. | low gamma | Fz-Cz | Latency | 0,975742021 | Not Significant |
| 40. | high gamma | Fz-Cz | Latency | 0,975742021 | Not Significant |
| 41. | low delta | Fz-Pz | Latency | 0,975742021 | Not Significant |
| 42. | high delta | Fz-Pz | Latency | 0,987853791 | Not Significant |
| 43. | low theta | Fz-Pz | Latency | 0,991378059 | Not Significant |
| 44. | high theta | Fz-Pz | Latency | 0,975742021 | Not Significant |
| 45. | low alpha | Fz-Pz | Latency | 0,975742021 | Not Significant |
| 46. | high alpha | Fz-Pz | Latency | 0,975742021 | Not Significant |
| 47. | low beta | Fz-Pz | Latency | 0,999432384 | Not Significant |
| 48. | high beta | Fz-Pz | Latency | 0,975742021 | Not Significant |
| 49. | low gamma | Fz-Pz | Latency | 0,975742021 | Not Significant |
| 50. | high gamma | Fz-Pz | Latency | 0,975742021 | Not Significant |
| 51. | low delta | Cz-Pz | Latency | 0,975742021 | Not Significant |
| 52. | high delta | Cz-Pz | Latency | 0,975742021 | Not Significant |
| 53. | low theta | Cz-Pz | Latency | 0,975742021 | Not Significant |

| | | | | | |
|---|---|---|---|---|---|
| 54. | high theta | Cz-Pz | Latency | 0,975742021 | Not Significant |
| 55. | low alpha | Cz-Pz | Latency | 0,991378059 | Not Significant |
| 56. | high alpha | Cz-Pz | Latency | 0,975742021 | Not Significant |
| 57. | low beta | Cz-Pz | Latency | 0,975742021 | Not Significant |
| 58. | high beta | Cz-Pz | Latency | 0,975742021 | Not Significant |
| 59. | low gamma | Cz-Pz | Latency | 0,462612914 | Not Significant |
| 60. | high gamma | Cz-Pz | Latency | 0,999432384 | Not Significant |
| 61. | low delta | Fp1-Fz | Duration | 0,462612914 | Not Significant |
| 62. | high delta | Fp1-Fz | Duration | 0,462612914 | Not Significant |
| 63. | low theta | Fp1-Fz | Duration | 0,975742021 | Not Significant |
| 64. | high theta | Fp1-Fz | Duration | 0,975742021 | Not Significant |
| 65. | low alpha | Fp1-Fz | Duration | 0,975742021 | Not Significant |
| 66. | high alpha | Fp1-Fz | Duration | 0,975742021 | Not Significant |
| 67. | low beta | Fp1-Fz | Duration | 0,975742021 | Not Significant |
| 68. | high beta | Fp1-Fz | Duration | 0,975742021 | Not Significant |
| 69. | low gamma | Fp1-Fz | Duration | 0,999432384 | Not Significant |
| 70. | high gamma | Fp1-Fz | Duration | 0,975742021 | Not Significant |
| 71. | low delta | Fp1-Cz | Duration | 0,975742021 | Not Significant |
| 72. | high delta | Fp1-Cz | Duration | 0,975742021 | Not Significant |
| 73. | low theta | Fp1-Cz | Duration | 0,975742021 | Not Significant |
| 74. | high theta | Fp1-Cz | Duration | 0,975742021 | Not Significant |
| 75. | low alpha | Fp1-Cz | Duration | 0,991378059 | Not Significant |
| 76. | high alpha | Fp1-Cz | Duration | 0,975742021 | Not Significant |
| 77. | low beta | Fp1-Cz | Duration | 0,991378059 | Not Significant |
| 78. | high beta | Fp1-Cz | Duration | 0,975742021 | Not Significant |
| 79. | low gamma | Fp1-Cz | Duration | 0,991378059 | Not Significant |
| 80. | high gamma | Fp1-Cz | Duration | 0,975742021 | Not Significant |
| 81. | low delta | Fp1-Pz | Duration | 0,975742021 | Not Significant |
| 82. | high delta | Fp1-Pz | Duration | 0,975742021 | Not Significant |
| 83. | low theta | Fp1-Pz | Duration | 0,975742021 | Not Significant |
| 84. | high theta | Fp1-Pz | Duration | 0,975742021 | Not Significant |
| 85. | low alpha | Fp1-Pz | Duration | 0,975742021 | Not Significant |
| 86. | high alpha | Fp1-Pz | Duration | 0,975742021 | Not Significant |
| 87. | low beta | Fp1-Pz | Duration | 0,462612914 | Not Significant |
| 88. | high beta | Fp1-Pz | Duration | 0,975742021 | Not Significant |
| 89. | low gamma | Fp1-Pz | Duration | 0,975742021 | Not Significant |
| 90. | high gamma | Fp1-Pz | Duration | 0,975742021 | Not Significant |
| 91. | low delta | Fz-Cz | Duration | 0,975742021 | Not Significant |
| 92. | high delta | Fz-Cz | Duration | 0,999432384 | Not Significant |
| 93. | low theta | Fz-Cz | Duration | 0,991378059 | Not Significant |
| 94. | high theta | Fz-Cz | Duration | 0,991378059 | Not Significant |
| 95. | low alpha | Fz-Cz | Duration | 0,987853791 | Not Significant |
| 96. | high alpha | Fz-Cz | Duration | 0,999432384 | Not Significant |
| 97. | low beta | Fz-Cz | Duration | 0,975742021 | Not Significant |
| 98. | high beta | Fz-Cz | Duration | 0,975742021 | Not Significant |
| 99. | low gamma | Fz-Cz | Duration | 0,975742021 | Not Significant |
| 100. | high gamma | Fz-Cz | Duration | 0,975742021 | Not Significant |
| 101. | low delta | Fz-Pz | Duration | 0,975742021 | Not Significant |
| 102. | high delta | Fz-Pz | Duration | 0,991378059 | Not Significant |
| 103. | low theta | Fz-Pz | Duration | 0,975742021 | Not Significant |
| 104. | high theta | Fz-Pz | Duration | 0,975742021 | Not Significant |
| 105. | low alpha | Fz-Pz | Duration | 0,975742021 | Not Significant |
| 106. | high alpha | Fz-Pz | Duration | 0,975742021 | Not Significant |
| 107. | low beta | Fz-Pz | Duration | 0,987853791 | Not Significant |
| 108. | high beta | Fz-Pz | Duration | 0,975742021 | Not Significant |
| 109. | low gamma | Fz-Pz | Duration | 0,975742021 | Not Significant |

| | | | | | |
|---|---|---|---|---|---|
| 110. | high gamma | Fz-Pz | Duration | 0,975742021 | Not Significant |
| 111. | low delta | Cz-Pz | Duration | 0,975742021 | Not Significant |
| 112. | high delta | Cz-Pz | Duration | 0,975742021 | Not Significant |
| 113. | low theta | Cz-Pz | Duration | 0,975742021 | Not Significant |
| 114. | high theta | Cz-Pz | Duration | 0,975742021 | Not Significant |
| 115. | low alpha | Cz-Pz | Duration | 0,462612914 | Not Significant |
| 116. | high alpha | Cz-Pz | Duration | 0,975742021 | Not Significant |
| 117. | low beta | Cz-Pz | Duration | 0,462612914 | Not Significant |
| 118. | high beta | Cz-Pz | Duration | 0,975742021 | Not Significant |
| 119. | low gamma | Cz-Pz | Duration | 0,975742021 | Not Significant |
| 120. | high gamma | Cz-Pz | Duration | 0,975742021 | Not Significant |
| 121. | low delta | Fp1-Fz | Mean Synchronization | 0,462612914 | Not Significant |
| 122. | high delta | Fp1-Fz | Mean Synchronization | 0,975742021 | Not Significant |
| 123. | low theta | Fp1-Fz | Mean Synchronization | 0,975742021 | Not Significant |
| 124. | high theta | Fp1-Fz | Mean Synchronization | 0,975742021 | Not Significant |
| 125. | low alpha | Fp1-Fz | Mean Synchronization | 0,975742021 | Not Significant |
| 126. | high alpha | Fp1-Fz | Mean Synchronization | 0,975742021 | Not Significant |
| 127. | low beta | Fp1-Fz | Mean Synchronization | 0,975742021 | Not Significant |
| 128. | high beta | Fp1-Fz | Mean Synchronization | 0,999432384 | Not Significant |
| 129. | low gamma | Fp1-Fz | Mean Synchronization | 0,975742021 | Not Significant |
| 130. | high gamma | Fp1-Fz | Mean Synchronization | 0,991378059 | Not Significant |
| 131. | low delta | Fp1-Cz | Mean Synchronization | 0,975742021 | Not Significant |
| 132. | high delta | Fp1-Cz | Mean Synchronization | 0,999432384 | Not Significant |
| 133. | low theta | Fp1-Cz | Mean Synchronization | 0,991378059 | Not Significant |
| 134. | high theta | Fp1-Cz | Mean Synchronization | 0,975742021 | Not Significant |
| 135. | low alpha | Fp1-Cz | Mean Synchronization | 0,987853791 | Not Significant |
| 136. | high alpha | Fp1-Cz | Mean Synchronization | 0,975742021 | Not Significant |
| 137. | low beta | Fp1-Cz | Mean Synchronization | 0,462612914 | Not Significant |
| 138. | high beta | Fp1-Cz | Mean Synchronization | 0,991378059 | Not Significant |
| 139. | low gamma | Fp1-Cz | Mean Synchronization | 0,975742021 | Not Significant |
| 140. | high gamma | Fp1-Cz | Mean Synchronization | 0,991378059 | Not Significant |
| 141. | low delta | Fp1-Pz | Mean Synchronization | 0,975742021 | Not Significant |
| 142. | high delta | Fp1-Pz | Mean Synchronization | 0,975742021 | Not Significant |
| 143. | low theta | Fp1-Pz | Mean Synchronization | 0,991378059 | Not Significant |
| 144. | high theta | Fp1-Pz | Mean Synchronization | 0,975742021 | Not Significant |
| 145. | low alpha | Fp1-Pz | Mean Synchronization | 0,975742021 | Not Significant |
| 146. | high alpha | Fp1-Pz | Mean Synchronization | 0,975742021 | Not Significant |
| 147. | low beta | Fp1-Pz | Mean Synchronization | 0,999432384 | Not Significant |
| 148. | high beta | Fp1-Pz | Mean Synchronization | 0,975742021 | Not Significant |
| 149. | low gamma | Fp1-Pz | Mean Synchronization | 0,975742021 | Not Significant |
| 150. | high gamma | Fp1-Pz | Mean Synchronization | 0,975742021 | Not Significant |
| 151. | low delta | Fz-Cz | Mean Synchronization | 0,999432384 | Not Significant |
| 152. | high delta | Fz-Cz | Mean Synchronization | 0,999432384 | Not Significant |
| 153. | low theta | Fz-Cz | Mean Synchronization | 0,975742021 | Not Significant |
| 154. | high theta | Fz-Cz | Mean Synchronization | 0,987853791 | Not Significant |
| 155. | low alpha | Fz-Cz | Mean Synchronization | 0,987853791 | Not Significant |
| 156. | high alpha | Fz-Cz | Mean Synchronization | 0,991378059 | Not Significant |
| 157. | low beta | Fz-Cz | Mean Synchronization | 0,975742021 | Not Significant |
| 158. | high beta | Fz-Cz | Mean Synchronization | 0,975742021 | Not Significant |
| 159. | low gamma | Fz-Cz | Mean Synchronization | 0,975742021 | Not Significant |
| 160. | high gamma | Fz-Cz | Mean Synchronization | 0,975742021 | Not Significant |
| 161. | low delta | Fz-Pz | Mean Synchronization | 0,975742021 | Not Significant |
| 162. | high delta | Fz-Pz | Mean Synchronization | 0,987853791 | Not Significant |
| 163. | low theta | Fz-Pz | Mean Synchronization | 0,975742021 | Not Significant |
| 164. | high theta | Fz-Pz | Mean Synchronization | 0,975742021 | Not Significant |
| 165. | low alpha | Fz-Pz | Mean Synchronization | 0,975742021 | Not Significant |

| | | | | | |
|---|---|---|---|---|---|
| 166. | high alpha | Fz-Pz | Mean Synchronization | 0,975742021 | Not Significant |
| 167. | low beta | Fz-Pz | Mean Synchronization | 0,999432384 | Not Significant |
| 168. | high beta | Fz-Pz | Mean Synchronization | 0,998362565 | Not Significant |
| 169. | low gamma | Fz-Pz | Mean Synchronization | 0,975742021 | Not Significant |
| 170. | high gamma | Fz-Pz | Mean Synchronization | 0,975742021 | Not Significant |
| 171. | low delta | Cz-Pz | Mean Synchronization | 0,975742021 | Not Significant |
| 172. | high delta | Cz-Pz | Mean Synchronization | 0,998362565 | Not Significant |
| 173. | low theta | Cz-Pz | Mean Synchronization | 0,975742021 | Not Significant |
| 174. | high theta | Cz-Pz | Mean Synchronization | 0,999432384 | Not Significant |
| 175. | low alpha | Cz-Pz | Mean Synchronization | 0,975742021 | Not Significant |
| 176. | high alpha | Cz-Pz | Mean Synchronization | 0,999432384 | Not Significant |
| 177. | low beta | Cz-Pz | Mean Synchronization | 0,987853791 | Not Significant |
| 178. | high beta | Cz-Pz | Mean Synchronization | 0,462612914 | Not Significant |
| 179. | low gamma | Cz-Pz | Mean Synchronization | 0,975742021 | Not Significant |
| 180. | high gamma | Cz-Pz | Mean Synchronization | 0,975742021 | Not Significant |
